# Efficient generation of targeted large insertions in mouse embryos using 2C-HR-CRISPR

**DOI:** 10.1101/204339

**Authors:** Bin Gu, Eszter Posfai, Janet Rossant

## Abstract

Rapid and efficient generation of large fragment targeted knock-in mouse models is still a major hurdle in mouse genetics. Here we developed 2C-HR-CRISPR, a highly efficient gene editing method based on introducing CRISPR reagents into mouse embryos at the 2-cell stage, taking advantage of the likely increase in HR efficiency during the long G2 phase and open chromatin structure of the 2-cell embryo. With 2C-HR-CRISPR and a modified biotin-streptavidin approach to localize repair templates to target sites, we rapidly targeted 20 endogenous genes that are expressed in mouse blastocysts with fluorescent reporters and generated reporter mouse lines. We showcase the first live triple-color blastocyst with all three lineages differentially reported. Additionally, we demonstrated efficient double targeting, enabling rapid assessment of the auxin-inducible degradation system for probing protein function in mouse embryos. These methods open up exciting avenues for exploring cell fate decisions in the blastocyst and later stages of development. We also suggest that 2C-HR-CRISPR can be a better alternative to random transgenesis by ensuring transgene insertions at defined ‘safe harbor’ sites.

---

The development of endonuclease-mediated genome editing technologies, in particular CRISPR/Cas9, has revolutionized mouse genetics by opening up the possibility to bypass the long and labor intensive embryonic stem cell-based methods and achieve precise genome editing directly in zygote stage embryos^1^. CRISPR/Cas9-mediated genome editing has shown consistently high efficiency in generating indels, point mutations and small insertions in zygotes^2,3^, largely making use of the non-homologous end-joining (NHEJ) and homology directed repair (HDR) pathways. However, the efficiency of precise introduction of large fragments by homologous recombination (HR) is still not ideal^4,5^. Previous reports have suggested methods to increase large fragment knock-in efficiency in zygotes, by Cas9-RNP injection^6^, with templates activating micro-homology mediated end joining or long-homology mediated end joining pathways^4,7^, with long singlestranded DNA templates or by chemically manipulating DNA repair pathways^8,9^. However, only a limited number of successfully targeted loci have been reported to date^4^. Some methods produce imprecise junctions at editing sites^7^, while others are limited by practical considerations and the size of the DNA fragment that can be inserted (up to 2 kb)^8^.

Fluorescent reporter mouse lines, which allow for direct visualization and live imaging of molecular and cellular dynamics during developmental, regenerative and pathological processes are among the most commonly generated models using large fragment knock-in. The preimplantation mouse embryo is an ideal model to study basic developmental mechanisms by live imaging^10^. Two consecutive crucial cell fate decisions separate embryonic and extraembryonic lineages and define three cell types in a 4-day time frame^10^. Moreover, because of the highly accessible nature of preimplantation embryos, they serve as excellent models to develop and apply cutting edge imaging and gene manipulation techniques such as single molecule tracking^11^, optogenetics and induced protein degradation systems^12,13^. Considering these advantages, we set out to generate a series of reporter mouse lines to study preimplantation development in embryos. The inconsistent and relatively low efficiency of knock-in reporter allele generation became a major hurdle and necessitated the development of methods for improved large fragment insertion in the mouse germ line.

HR is predominantly active in late S/G2 phases of the cell cycle^14^. It has been shown that timely delivery of Cas9-RNP and donor DNA into G2-synchronised cells or restricting the presence of Cas9 protein to late S/G2 phases by fusing it with Geminin *in vitro* significantly increased knock-in efficiency^15–17^. Approaching the problem from a mouse developmental biologist’s perspective, we reasoned that the 2-cell stage might be an ideal window for introducing large DNA fragments by HR, as the G2 phase during this cell cycle is exceptionally long (10-12 hours) and is relatively synchronized between developing embryos. Additionally, major zygotic genome activation, which takes place during the extended G2 phase of the 2-cell stage, is associated with an open chromatin state and would thus likely increase the accessibility of the chromatin to editing enzymes and repair templates (Fig. 1a). We therefore set out to generate large fragment knock-in alleles in mouse embryos by injecting CRISPR/Cas9 reagents and repair templates into 2-cell embryos (2C-HR-CRISPR). We first compared the knock-in efficiencies at the 2-cell and zygote stages by generating two reporter alleles: a) the fusion of a minimum auxin degron (mAID) domain^18^ and mCherry to the C-terminus of Sox2 (Sox2-mAID-mCherry)(with 5 and 1 kb homology arms) and b) the fusion of the Halo-tag^19^ to the C-terminus of Gata6 (with 2 and 3 kb homology arms)(Fig. 1b). We microinjected *Cas9* mRNA, sgRNA and a circular donor plasmid into the cytoplasm of either zygotes or both cells of 2cell embryos (Supplementary Movie 1). As both genes have highly specific expression patterns in the blastocyst, we evaluated targeting efficiencies by culturing microinjected embryos to the blastocyst stage and scoring for the presence and localization of the fluorescent tag by both live imaging and co-immunostaining with lineage markers (Supplementary Fig. 1). The knock-in efficiency increased from less than 1% with zygote microinjection to 31.5% with 2cell microinjection for Sox2-mAID-mCherry and from 6.5% to 35% for Gata6-Halo (Fig. 1b). We noted mosaicism in targeted embryos (Supplementary Fig. 2), similar to observations in founder animals generated by CRISPR-editing in zygotes. This is likely due to varied independent editing of alleles in S/G2 phase zygotes (4 alleles for most genes) and G2 phase 2-cell embryos (8 alleles for most genes). While the increase in allele numbers at the 2-cell stage may partly contribute to the increased knock-in efficiency, CRISPR-mediated gene editing at 2-cell stage increased knock-in efficiency well beyond 2-fold, demonstrating a real increase in targeting efficiency per allele by 2-cell microinjection.

**Figure 1.**
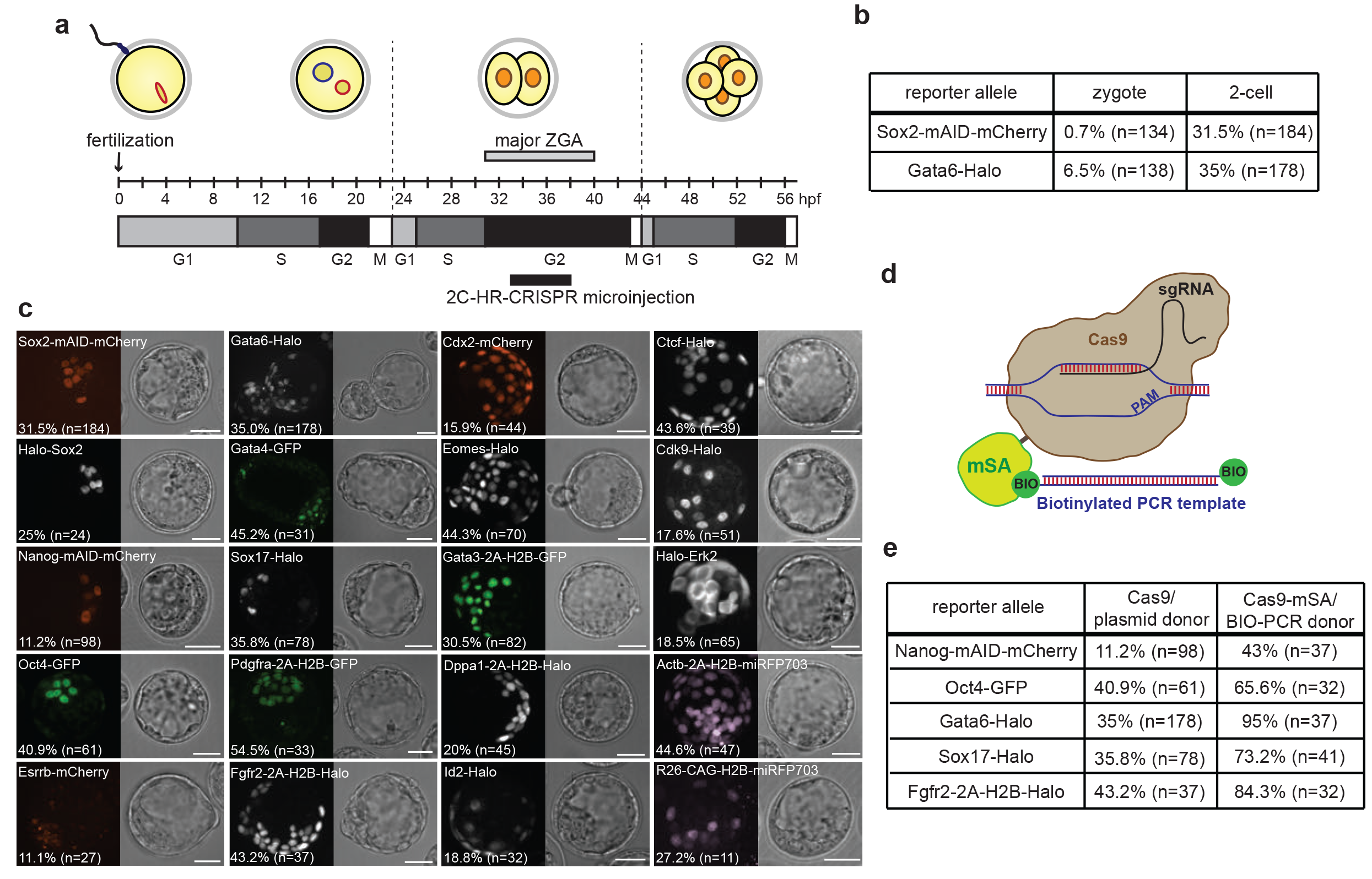
Efficient large fragment targeting using HR-mediated CRISPR/Cas9 editing in 2-cell stage mouse embryos (2C-HR-CRISPR). **(a)** Schematic showing timing of 2C-HR-CRISPR microinjections relative to cell cycle progression and ZGA in the mouse embryo. Hpf - hours post fertilization. **(b)** Efficiency of inserting large tags at the C-terminus of Sox2 and Gata6 coding sequences when microinjecting zygotes versus 2-cell stage embryos. Targeting efficiencies were evaluated by expression at the blastocyst stage. Sox2 expression is restricted to the epiblast (EPI) lineage, while Gata6 is expressed in the primitive endoderm (PE) and to some extent the trophectoderm (TE) lineages. n=number of microinjected embryos which developed to the blastocyst stage. c) The catalogue of pre-implantation markers tagged using 2C-HR-CRISPR. Live images of 2C-HR-CRISPR microinjected embryos cultured to the blastocyst stage. All embryos are likely mosaic, but contain at least one correctly targeted allele in some cells. The Halo tag was visualized using the live JF549, JF635 or JF646 dyes. Targeting efficiencies achieved at the blastocyst stage are indicated, n=number of microinjected embryos, which developed to the blastocyst stage. Bright field images are single plane; fluorescent images show extended focus. Scale bars: 40 μm. **(d)** Schematic showing the Cas9-mSA/BIO-PCR donor CRISPR system. **(e)** Comparison of targeting efficiencies of large tags at different loci using Cas9/plasmid donor or Cas9-mSA/BIO-PCR donor 2C-HR-CRISPR microinjections. Targeting efficiency was evaluated at the blastocyst stage, n=number of microinjected embryos, which developed to the blastocyst stage.

To demonstrate the wider applicability of the 2C-HR-CRISPR method, we generated reporter alleles (with up to 4.5 kb insert size) for 20 loci in the mouse genome (Fig. 1c). We mostly constructed C or N-terminal fusions or in some cases using the self-cleaving T2A peptide^20^ (T2A-H2B-reporter) with different reporter tags such as GFP, mCherry, Halo^19^ and miRFP703^21^. We selected most “classical” lineage markers covering all three cell types of the blastocyst. These included epibalst (EPI) (*Sox2*, *Nanog*, *Oct4* and *Esrrb*), primitive endoderm (PE) (*Gata6*, *Gata4*, *Sox17*, *Fgfr2* and *Pdgfra*) and trophectoderm (TE) (*Cdx2*, *Eomes* and *Gata3*) markers^10^ (Fig. 1c). We also targeted *Id2* and *Dppa1*, two genes which have so far only been predicted to be TE-specific lineage markers based on mRNA profiles^22,23^. Additionally, we selected a number of other genes, including the chromatin structural regulator *Ctcf*, the transcriptional elongation factor *Cdk9*, the cell signaling kinase *Mapk1* (*Erk2*) and two safe-harbor loci where insertion of transgenes provides high and uniform expression in all cells (the *Rosa26* (*R26*)^24^ and the *beta-Actin* (*Actb*) loci). Importantly, we succeeded in targeting all 20 selected loci, as read out by correct lineage and subcellular localization of the tag in microinjected embryos cultured to the blastocyst stage and validated correct targeting either by PCR genotyping single embryos or by immunostaining targeted blastocysts where antibodies against the endogenous protein were available (Supplementary Fig. 1). We found that targeting showed a highly reproducible locus and sgRNA-dependent efficiency (Fig. 1c and Supplementary Fig. 3). However, we did not find any loci that we could not target. Overall knock-in efficiency varied between 11-55% depending on the locus. Additionally, we showed that targeting efficiencies were comparable on an outbred or inbred (C57Bl/6J) mouse background (Supplementary Fig. 4). Therefore, we demonstrated that 2C-HR-CRISPR is a highly robust method to generate large fragment knock-in alleles.

Apart from the activity of HR pathway, the availability of the repair template could be another limiting factor for knock-in generation. Methods to recruit the repair template to the editing site have been explored to improve targeting efficiencies^25,26^. We therefore constructed a fusion gene between Cas9 and monomeric streptavidin (Cas9-mSA)^27^ and generated biotinylated repair templates containing homology arms and insertion fragments by PCR amplification (BIO-PCR) from targeting vectors with 5’-biotinylated primers (Fig. 1d). Using this Cas9-mSA/BIO-PCR method we further improved targeting efficiencies at each of the 5 genes tested (Fig. 1e), reaching an extreme of 95% knock-in efficiency when targeting a Halo-tag to the C-terminus of Gata6. We also observed an increase in the number of fluorescently labeled cells in individual embryos, suggesting an increase in the number of alleles correctly targeted and consequently a likely decrease in mosaicism. Moreover, from two out of the three male Gata6-Halo founder mice generated using the Cas9-mSA/BIO-PCR method (see later) we obtained 100% Gata6-Halo positive blastocysts by breeding to CD1 females suggesting that all germ cell progenitors were homozygously targeted.

We then transferred microinjected embryos to generate reporter mouse lines (Fig. 2a). These included three reporter lines for preimplantation cell lineages: *Sox2-mAID-mCherry* and *Nanog-mAID-mCherry* for the EPI lineage and *Gata6-Halo* for the PE lineage, a *Cdk9-Halo* line for imaging transcription dynamics in early embryos, and a far red pan-nuclear marker line *R26-CAG-H2B-miRFP703*. We successfully obtained positive founder animals from all transfer sessions and validated correct targeting by long-range genotyping PCR. Importantly, we obtained germ line transmission from all founders tested by both visualizing the correct localization of reporters in N1 generation blastocysts (Fig. 2b) and genotyping N1 animals. Furthermore, we validated the correct sequence of the knock-in allele in N1 animals by Sanger sequencing the 5’ and 3’ junctions of the insertion, including the homology arms (data not shown) and excluded the possibility of random integrations and concatemer insertions by digital droplet PRC. We did not rule out the possibility of off-target effects, however, as outcrossing is necessary to deplete incorrectly edited alleles, other byproducts of CRISPR-mediated editing will also be bred out during mouse line production and thus do not pose a threat. Moreover, as 2C-HR-CRISPR does not involve any manipulation of the cell cycle or DNA-repair pathways, as attempted in other previous studies, the likelihood of undesirable genomic perturbations is minimal.

**Figure 2.**
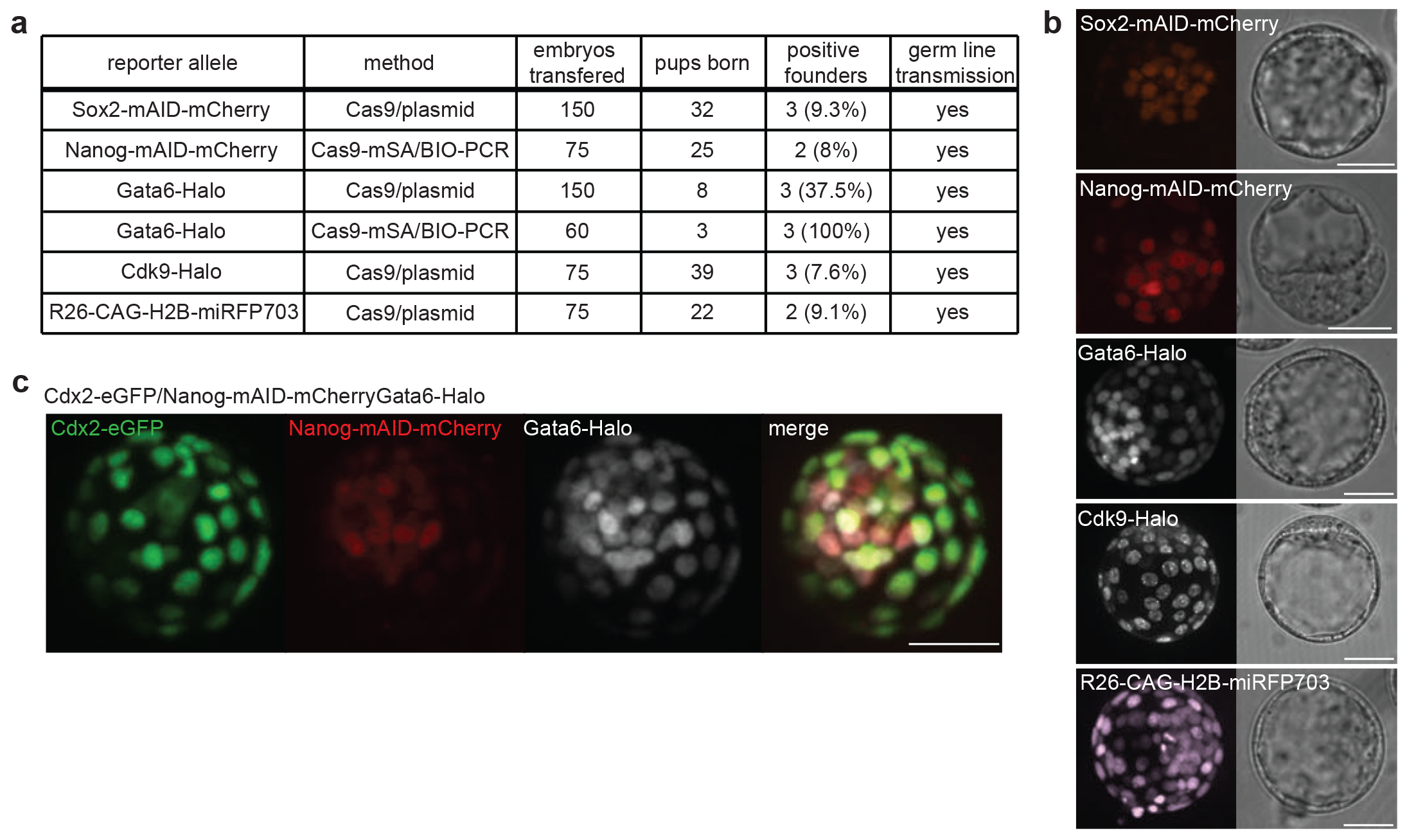
Generation of mouse lines using 2C-HR-CRISPR. **(a)** Table showing mouse lines generated using either the Cas9/plasmid donor or Cas9-mSA/BIO-PCR donor 2C-HR-CRISPR methods. **(b)** Live image of N1 generation blastocysts isolated from founder mice for all mouse lines generated. All cells in the embryo are heterozygous, non-mosaic for the targeted reporter allele. The Halo-tag was visualized using the live JF549 dye. Bright field images are single plane; fluorescent images show extended focus. Scale bars: 40 μm. **(c)** Non-mosaic live triple-color blastocyst with all three lineages labeled with different reporter alleles. Embryo was generated by breeding 2C-HR-CRISPR-generated mouse lines (*Nanog-mAID-mCherry* and *Gata6-Halo*) with an existing *Cdx2-eGFP* reporter mouse line. The Halo-tag was visualized using the live JF646 dye. Bright field image is single plane; fluorescent images show extended focus. Scale bar: 40 μm.

By breeding 2C-HR-CRISPR-generated mouse lines (*Nanog-mAID-mCherry* and *Gata6-Halo*) to an existing *Cdx2-eGFP* reporter line^28^ we were able to produce the first live triple-color blastocysts, in which all cells were marked by a reporter in a lineage dependent manner (Fig. 2c). Multi-color reporter embryos will be invaluable tools for exploring lineage segregation in real time by live imaging and resolving long-standing controversies stemming from attempts to reconstruct dynamic developmental processes from static snapshots. As proof of principle, we followed the development of *Cdx2-eGFP/Gata6-Halo* embryos (1 Z-stack/hour) for 30 hours from the 16-cell to the blastocyst stage by live imaging (Supplementary Fig. 5 and Supplementary Movie 2). However, we noted that phototoxicity became a limiting factor in acquiring multi-color images with adequately high spatial and temporal resolution. Recent advances in high-resolution light sheet imaging could overcome these limitations^29,30^ and thus are expected to significantly advance the applicability of multi-color reporter lines. We also anticipate our reporter mouse lines to be valuable resources beyond the preimplantation field, as many of these genes are expressed in multiple other cell types where their roles can be interrogated by live imaging.

Given the high frequency of targeting by 2C-HR-CRISPR, we attempted to achieve double targeting in embryos by simultaneously tagging *Nanog* with mAID-mCherry using the Cas9-mSA/BIO-PCR method and *Gata6* with the Halo-Tag using the Cas9/plasmid method. Efficient (29%) precise insertion of both large fragments in the same embryo was achieved (Fig. 3a and b). We then tested the delivery and the applicability of the auxin-inducible degron system in embryos, a two-component tool for inducible, rapid and reversible protein degradation (Fig. 3c), which has so far only been utilized in yeast^13^, *C. elegans*^*31*^ or mammalian cell culture systems^18^.

**Figure 3.**
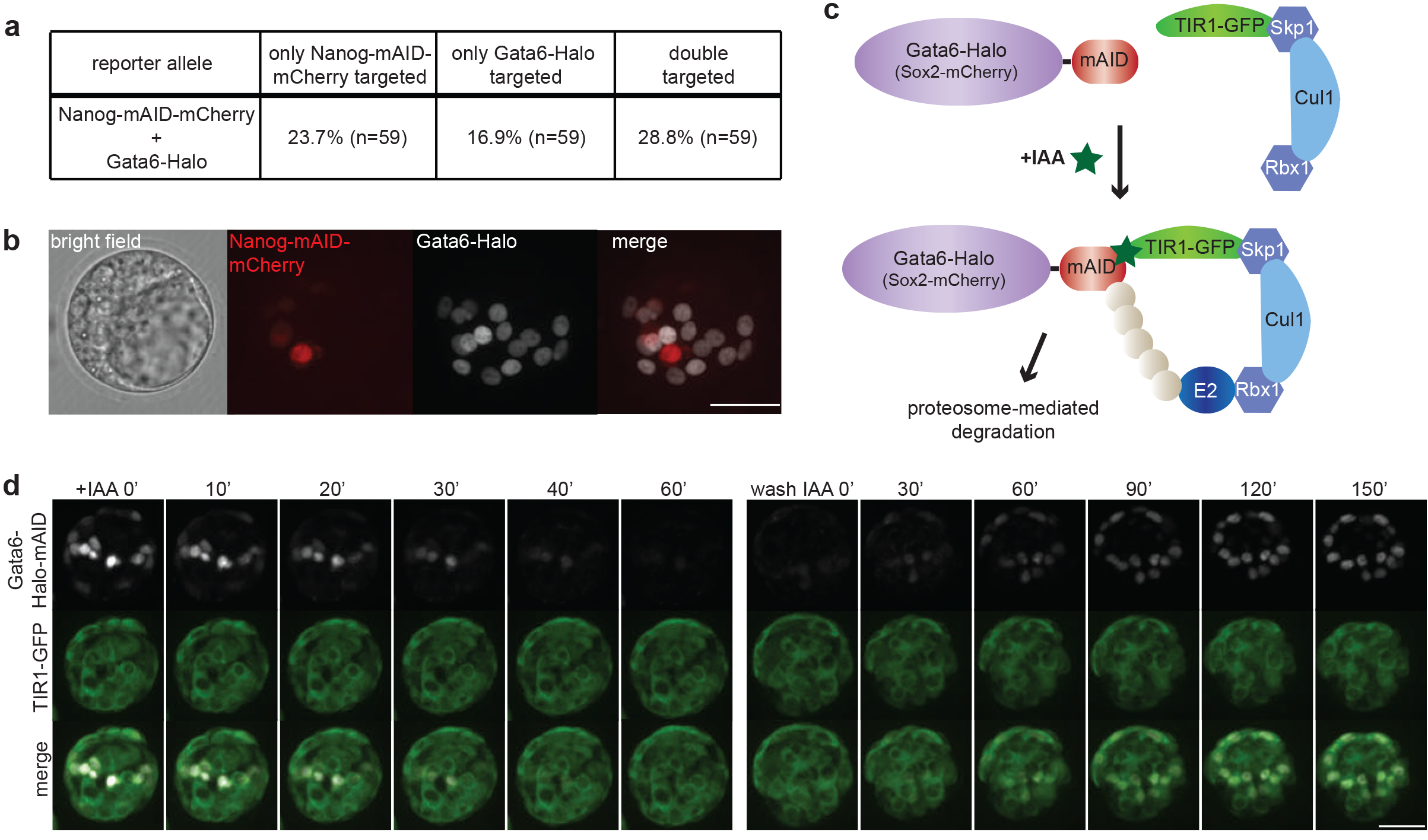
Simultaneous targeting of two different large inserts and the establishment of the auxin-inducible degron system in the mouse embryo. **(a)** Targeting efficiencies in double-targeting experiments. Embryos were co-microinjected with reagents to generate *Gata6-Halo* (Cas9/plasmid donor) and *Nanog-mAID-mCherry* (Cas9-mSA/BIO-PCR donor) at the 2-cell stage. Targeting efficiencies were evaluated at the blastocyst stage, n=number of microinjected embryos, which developed to the blastocyst stage. **(b)** Example live image of double-targeted embryo cultured to the blastocyst stage. The Halo-tag was visualized using the live JF646 dye. Bright field image is single plane; fluorescent images show extended focus. Embryo is mosaic for Gata6-Halo and Nanog-mAID-mCherry. Scale bars: 40 μm. **(c)** Schematic of the auxin-inducible degron system used in preimplantation embryos. **(d)** Time-lapse imaging of auxin-induced degradation of Gata6-Halo-mAID in a TIR1-GFP positive embryo. Simultaneous targeting of Halo-mAID to the *Gata6* locus and TIR1-GFP to the *Actb* locus was achieved with 2C-HR-CRISPR. Left panel shows embryo at different times after addition of auxin (+IAA) and right panel shows same embryo at different times following washing out IAA. All images show extended focus. Scale bar: 40 μm.

We simultaneously targeted *TIR1-GFP* to the *Actb* locus (*Actb-T2A-TIR1-GFP*) and Halo-mAID to the C-terminus of *Gata6* (*Gata6-Halo-mAID*) or mAID-mCherry to the C-terminus of *Sox2* and identified blastocysts with double-targeted cells. TIR1, a plant-specific F-box protein recruits the SCF ubiquitin ligase complex^18^ to mAID-tagged proteins in an auxin (IAA)-dependent manner, which results in their polyubiquitination and subsequent proteasome-mediated degradation. Following addition of IAA to embryos with double-targeted cells we found that in TIR1-GFP positive cells, Gata6-Halo-mAID was depleted to background levels in 30-60 minutes and Sox2-mAID-mCherry was degraded in 30-80 min, while in TIR1-GFP negative cells expression of the reporters remained stable (Fig 3d and Supplementary Fig. 6). We then washed out IAA and traced the recovery of proteins for 2-3 hours. We found that both Gata6-Halo-mAID and Sox2-mAID-mCherry gradually re-appeared in TIR1-GFP positive cells within this time frame. Additionally, we confirmed that culturing preimplantation embryos in the presence of IAA had no adverse effect on development (data not shown). We thus demonstrate for the first time the use of the auxin-inducible degron system in mouse embryos for inducible and reversible degradation of endogenous proteins.

In summary, 2C-HR-CRISPR is a robust method to insert any large fragment in a site-specific manner into the mouse genome, accelerating transgenic knock-in mouse production. In addition to targeting endogenous genes, 2C-HR-CRISPR can be used to insert transgenes at defined ‘safe harbor’ sites, thus avoiding position effects that accompany standard trangenesis and can facilitate the application of synthetic biology tools in mouse models.

## Acknowledgements

We thank the Transgenic Core lab headed by Marina Gertsenstein at the Centre for Phenogenomics, Toronto for transgenic services and useful discussions, Dr Daniel Durocher for discussion on the parameters affecting HR, and Dr. Yojiro Yamanaka for previously setting up the 2-cell microinjection system in the Rossant lab. This work was funded by CIHR (FDN-143334) and Genome Canada and Ontario Genomics (OGI-099).

### Author contributions

B.G., E.P. and J.R. conceived the study. B.G. had the idea of increasing HR by microinjecting 2-cell embryos. B.G. and E.P. designed, carried out and analyzed all experiments equally. J.R. provided supervision and funding for the study. All authors contributed to writing the manuscript.

### Competing Financial Interest

Authors declare no competing financial interests.

## Methods

### Mouse lines and embryos

CD1(ICR) (Charles River), C57Bl/6J (Charles River) and *Cdx2-eGFP* (knock-in fusion to endogenous locus)^1^ mouse lines were used as embryo donors in this study. Embryos were collected from superovulated and mated females as described before^2^. Zygotes were isolated from the oviduct at 0.5 days post coitum (dpc) and washed clean of cumulus cells by brief treatment with 300μg/ml hyaluronidase (Sigma). 2-cell embryos were collected at 1.5 dpc and blastocysts at 3.5 dpc. If not immediately used, embryos were cultured in small drops of KSOM supplemented with amino acids (EMD Milipore) under mineral oil (Zenith Biotech) at 37°C, with 6% CO_2_ for specified times. Founder mice were genotyped by PCR amplification with primers spanning homology arms. Founder mice were out-crossed to CD1 mice to generate N1 mice. The N1 mice were genotyped by PCR. Additionally, genomic regions spanning the targeting cassette and 3’ and 5’ homology arms were Sanger-sequenced to validate correct targeting and insertion copy number was evaluated by droplet digital PCR (performed by the Centre for Applied Genomics at the Research Institute of The Hospital for Sick Children, Toronto). All animal work was carried out following Canadian Council on Animal Care Guidelines for Use of Animals in Research and Laboratory Animal Care under protocols approved by The Centre for Phenogenomics Animal Care Committee (20-0026H).

### Oligos

DNA oligos were synthesized by either Sigma or IDT and all sequences used in this study are provided in Supplementary Table 1.

### Constructs

To construct pCS2+-Cas9-mSA, the mSA coding sequence was amplified from pDisplay-mSA-EGFP-TM (a gift from Dr. Sheldon Park (Addgene plasmid #39863))^3^ and fused to the C-terminus of the Cas9 coding sequence with an optimized linker^4^ in pCS2+-Cas9 using InFusion cloning. For the construction of EasyFusion plasmids, the mCherry cassette in the pBluescript-mCherry vector was replaced with different insertion cassettes by InFusion cloning^5–7^. For constructing donor vectors, homology arms of 1-5 kb length were amplified from BACs covering the genome sequence of interest and inserted into EasyFusion vectors by InFusion cloning. For the Rosa-CAG-H2B-miRFP703 vector H2B-miRFP703 was inserted into the MluI site of R26-CAG-Asisl:MluI plasmid (a gift from Dr. Ralf Kuehn (Addgene plasmid # 74286))^8^ and the floxed stop cassette was removed by *in vitro* Cre reaction. sgRNA target sequences were selected around the start (for N-terminal targeting) or stop (for C-terminal targeting) codon (for exact design see Supplementary Fig. 3) and off target sites were evaluated using the Optimized CRISPR design tool (http://crispr.mit.edu). sgRNAs with predicted off-target sites in coding regions were excluded. sgRNA cloning into pX330 vectors (a kind gift from Dr. Feng Zhang (Addgene plasmid #42230)) was performed as previously described^9^.

### EasyFusion donor vectors for rapid cloning of knock-in targeting constructs

To further simplify the process of targeting vector construction and facilitate the workflow of 2C-HR-CRISPR methods, we constructed EasyFusion vectors. This set of general vectors consists of a module of interest for knock-in (e.g. fluorescent proteins (FP) or the Halo-tag compatible with C- or N-terminal fusions; 2A-FPs/Halo for C-terminal targeting or the mAID degron domain for inducible protein degradation), flanked with multiple unique restriction sites available for insertion of homology arms using inFusion cloning. All EasyFusion constructs will be available from Addgene.

### RNA synthesis

RNAs were synthesized as described before^2^. Briefly, to produce *Cas9* or *Cas9-mSA* mRNA, the pCS2+-Cas9 or pCS2+-Cas9-mSA plasmid was linearized with NotI restriction digestion and used as template for *in vitro* transcription using the mMESSAGE mMACHINE^®^ SP6 Transcription Kit (Thermo Fisher Scientific). For the production of sgRNAs, sgRNA coding sequences were PCR-amplified from the pX330 plasmids with primers containing the T7 promoter and used as template to produce sgRNA using the MEGAshortscript^™^ T7 Transcription Kit (Thermo Fisher Scientific). All RNA products were purified using the RNeasy Mini Kit (Qiagen) following the clean-up protocol.

### Cytoplasmic microinjection of zygote and 2-cell stage embryos

Cytoplasmic microinjection of zygotes and 2-cell stage embryos were performed as described previously^10^. Briefly, *Cas9* mRNA (100ng/μl) or *Cas9-mSA* mRNA (75ng/μl), sgRNA (50ng/μl) and donor plasmids (30ng/μl) or BIO-PCR donors (20ng/μl) were microinjected in nuclease-free injection buffer (10mM Tris-HCl, pH 7.4, 0.25mM EDTA). Microinjection was performed using a Leica microscope and micromanipulators (Leica Microsystems Inc). Injection pressure was provided by a FemtoJet (Eppendorf) and negative capacitance was generated using a Cyto721 intracellular amplifier (World Precision Instruments). Microinjections were performed in M2 medium (Zenith Biotech) in an open glass chamber or under mineral oil. Following microinjections, embryos were cultured in KSOM either to the morula stage (8 to 16-cell) for embryo transfers or to blastocyst stage for evaluating targeting efficiencies. To generate mouse lines, 25-30 microinjected embryos were transferred into the oviducts of pseudopregnant females one day after microinjections. To evaluate targeting efficiencies embryos were cultured to the blastocyst stage. The Halo tag was visualized using 10nM JF549, JF635 or JF646 dyes (a gift from Dr. Luke Lavis)^11^.

### Whole-mount immunofluorescent staining of embryos

Whole mount immunofluorescent staining of embryos was performed as previously described^10^. Briefly, embryos were fixed in 4% paraformaldehyde at room temperature for 15 minutes, washed once in PBS containing 0.1% Triton X-100 (PBS-T), permeabilized for 15 minutes in PBS 0.5% Triton X-100 and then blocked in PBS-T with 2% BSA (Sigma) and 5% normal donkey serum (Jackson ImmunoResearch Laboratories) at room temperature for 2 hours, or overnight at 4°C. Primary and secondary antibodies were diluted in blocking solution. Staining was performed at room temperature for ~2 hours or overnight at 4°C. Washes after primary and secondary antibodies were done three times in PBS-T. Embryos were mounted in Vectashield containing Dapi (Vector Laboratories) in wells made with Secure Seal spacers (Molecular Probes™, Thermo Fisher Scientific) and placed between two cover glasses for imaging. Primary antibodies: chicken anti-mCherry 1:600 (NBP2-25158, Novus Biologicals); mouse anti-mCherry 1:500 (632543, Clontech); chicken anti-GFP 1:400 (ab13970, Abcam); rabbit anti-Cdx2 1:600 (ab76541, Abcam); mouse anti-Cdx2 1:100 (MU392-UC, Biogenex); rabbit anti-Eomes 1:200 (23345, Abcam); mouse anti-Nanog 1:200 (560259, BD Biosciences); goat anti-Sox2 1:100 (AF2018, RandD Systems); goat anti-Oct4 1:100 (sc-8629, Santa Cruz Biotechnologies); mouse anti-Esrrb 1:100 (PP-H6705-00, Perseus Proteomics); goat anti-Sox17 1:100 (AF1924, RandD Systems); rabbit anti-Gata4 1:100 (sc-9053, Santa Cruz Biotechnologies) and goat anti-Gata6 1:100 (AF1700, RandD Systems). The Halo tag was not stained using an antibody, as the JF549 dye remained fluorescent following fixation. Secondary antibodies: (diluted 1:500) 448, 549 or 633 conjugated donkey anti-mouse, donkey anti-rabbit or donkey anti-goat DyLight (Jackson ImmunoResearch) or Alexa Fluor (Life Technologies).

### Single blastocyst genotyping

Single blastocyst genotyping was performed as previously described^12^. Briefly, microinjected embryos cultured to the blastocyst stage were imaged on a spinning disk confocal microscope to evaluate targeting. Genomic DNA of single blastocysts was extracted with Red Extract-N-Amp kit (Sigma) and PCR genotyping was performed with gene-specific primers using PhireII polymerase (Thermo Fisher).

### Confocal microscopy and image analysis

Both live and immunostained images were acquired using a Zeiss Axiovert 200 inverted microscope equipped with a Hamamatsu C9100-13 EM-CCD camera, a Quorum spinning disk confocal scan head and Volocity acquisition software version 6.3.1. Single plane images or Z-stacks (at 1μm intervals) were acquired with a 40x air (NA=0.6) or a 20x air (NA=0.7) objective. Images were analyzed using Volocity.

Time-lapse imaging was performed on the same microscope equipped with an environment controller (Chamlide, Live Cell Instrument, Seoul, South Korea). Embryos were placed in a glass-bottom dish (MatTek) in KSOM or phenol-red free mouse embryonic stem cell medium (Thermo Fisher) (for blastocyst stage embryos) covered with mineral oil. A 20x air (NA=0.7) objective lens was used. Images were acquired at 3μm Z intervals with time-lapse settings as indicated in figure legends.

### Auxin-inducible protein degradation

Embryos were simultaneously targeted with Actb-T2A-TIR1-eGFP and Gata6-Halo-mAID or Sox2-mAID-mCherry constructs using the combination of Cas9/plasmid donor and Cas9-mSA/BIO-PCR donor 2C-HR-CRISPR methods. Experimental embryos with double positive cells or control embryos with only Gata6-Halo-mAID or Sox2-mAID-mCherry expressing cells were identified at the blastocyst stage. Protein degradation was induced by addition of 500μM IAA (Sigma) to the culture media, and degradation was stopped by washing embryos through several drops of IAA-free media. Degradation and protein recovery was recorded with time-lapse imaging for times indicated in figures.

### Statistical analysis

No statistical methods were used to pre-determine sample size. The experiments were not blinded from investigators: the identities of the samples were known throughout the experiment. No statistical tests were performed in this study.

---

**Supplementary Figure 1.**
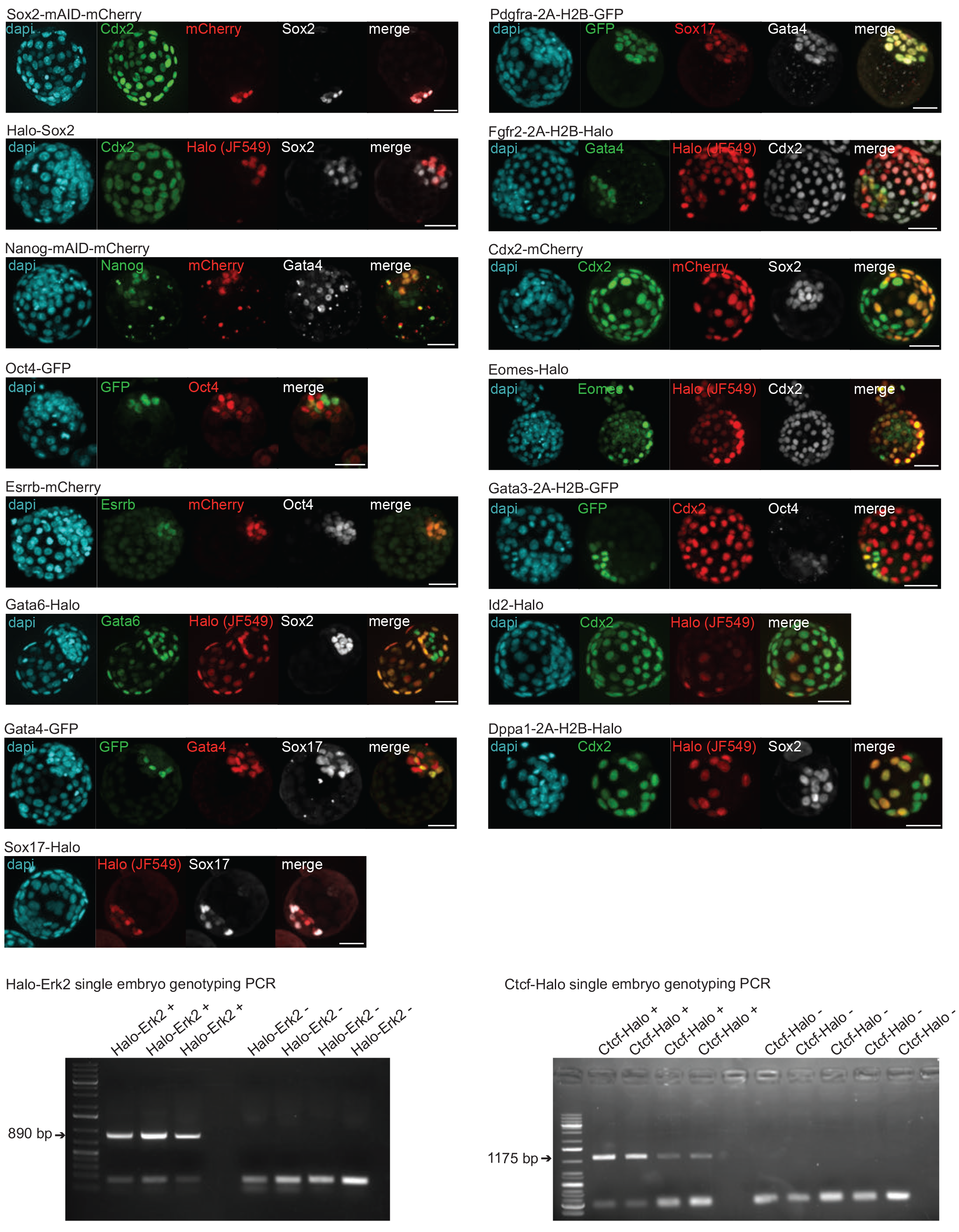
Validation of correct targeting at each locus by immunofluorescent staining of 2C-HR-CRISPR microinjected embryos cultured to the blastocyst stage. Embryos were stained for the inserted tag and targeted endogenous gene product (Sox2, Nanog, Oct4, Esrrb, Klf4, Gata6, Gata4, Sox17, Cdx2, Eomes) or where antibodies for the endogenous protein were not available (Fgfr2, Pdgfra, Id2, Dppa1, Gata3) correct lineage localization of the tag was confirmed using established lineage markers. Extended focus images are shown. Merged image only contains tag and endogenous protein expression (Sox2, Nanog, Oct4, Esrrb, Klf4, Gata6, Gata4, Sox17, Cdx2, Eomes) or tag and appropriate lineage marker (Fgfr2, Pdgfra, Id2, Dppa1, Gata3). Scale bars: 40 μm. PCR genotyping gel confirming of Halo-Erk2 and Ctcf-Halo targeting by genotyping single targeted and non-targeted (negative control) blastocysts. Targeted and non-targeted embryos were pre-selected by checking for the presence of fluorescently labeled (JF549) Halo-tag at the blastocyst stage.

**Supplementary Figure 2.**
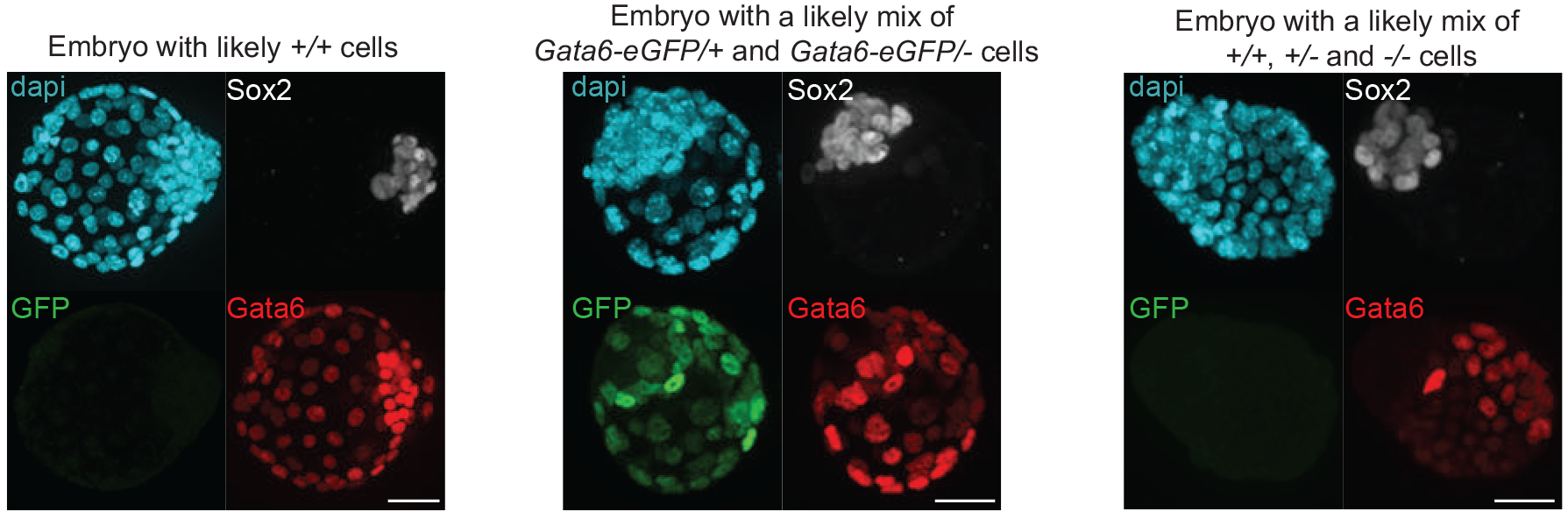
Embryos microinjected with CRISPR reagents targeting Gata6-GFP using 2C-HR-CRISPR. All embryos are from the same microinjection session. Embryos were cultured to the blastocyst stage and immunostained with indicated antibodies. Scale bars: 40 μm. Extended focus images are shown. Left embryo likely has wild-type/wild-type (*+/+*) cells, due to the uniform expression of endogenous Gata6 in PE (stronger) and TE (weaker) cells and no detectable GFP signal. Middle embryo likely has a mix of *Gata6-GFP/+* and *Gata6-GFP/deleted* (−) cells, as GFP is detected in seemingly all Gata6 positive cells, but endogenous Gata6 expression levels vary. Right embryo likely has a mix of *+/+*, *+/−* and *−/−* cells, corresponding to strong, intermediate and completely absent endogenous Gata6 signals.

**Supplementary Figure 3.**
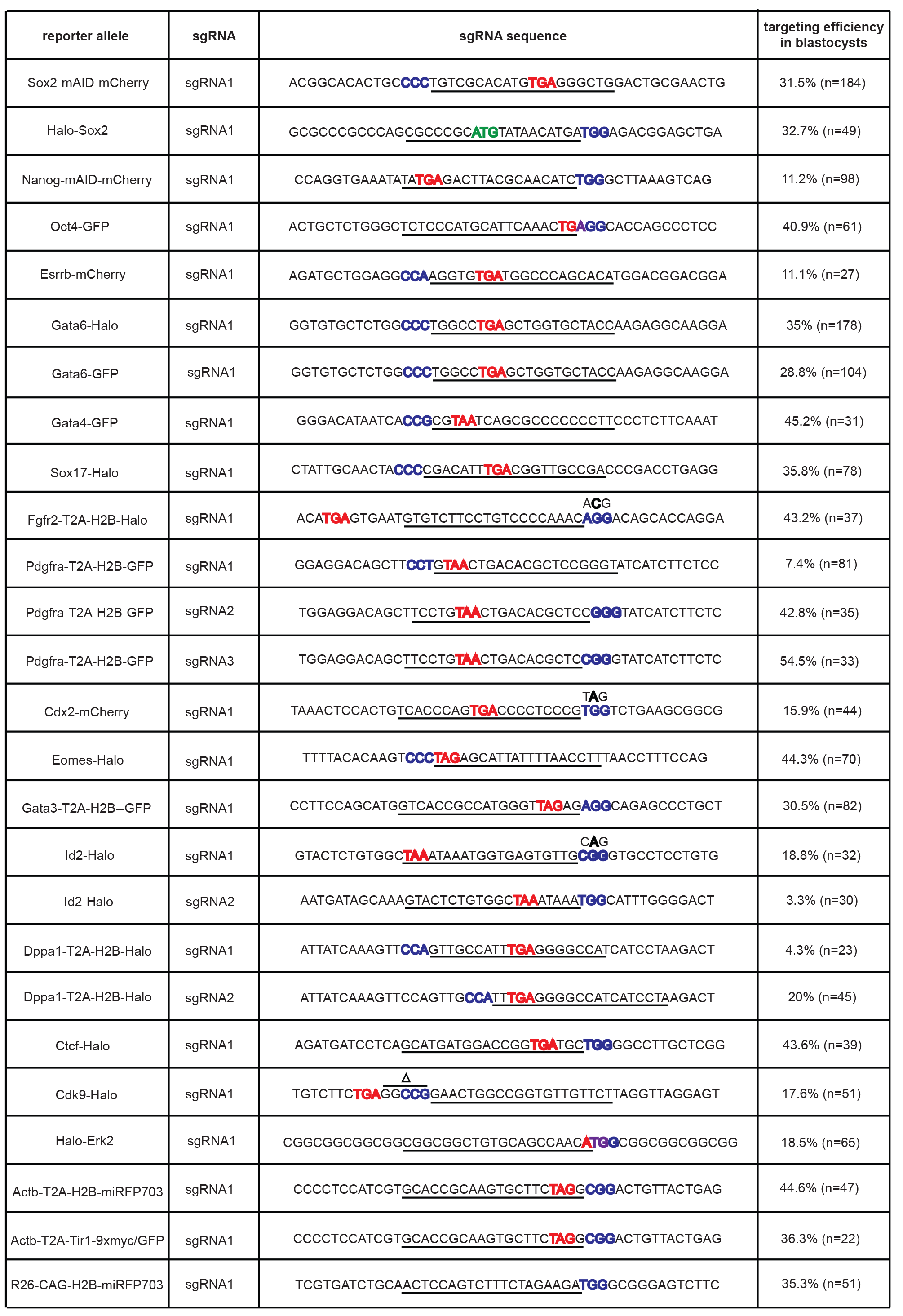
Table showing all loci targeted, sgRNAs used and targeting efficiencies achieved at the blastocyst stage using 2C-HR-CRISPR (n indicates the number of microinjected embryos, which developed to the blastocyst stage). sgRNA sequence is indicated in the genomic sequence surrounding the insertion site: the N_20_ bases of the sgRNA are underlined, the PAM sequence is in blue. Start or stop codons are highlighted in red or green, respectively. If a synonymous mutation was introduced into the targeting donor to avoid cleavage by the sgRNA/Cas9, this is indicated above the DNA sequence.

**Supplementary Figure 4.**
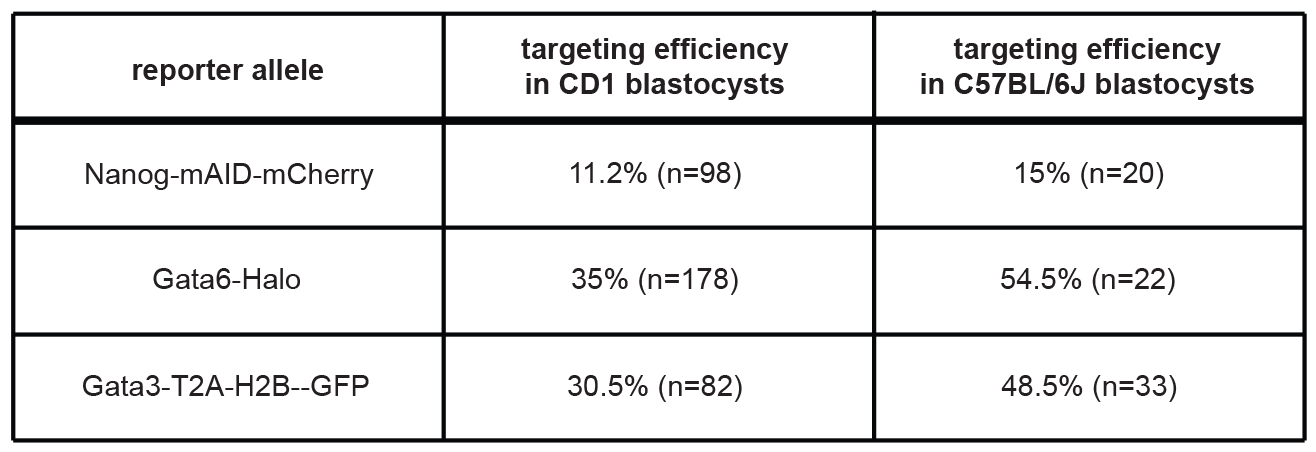
2C-HR-CRISPR targeting efficiencies compared between CD1 and C57BL/6J backgrounds. Microinjected embryos were cultured to the blastocyst stage where targeting efficiency was evaluated by scoring correct localization of reporter expression (n indicates number of microinjected embryos that developed to the blastocyst stage).

**Supplementary Figure 5.**
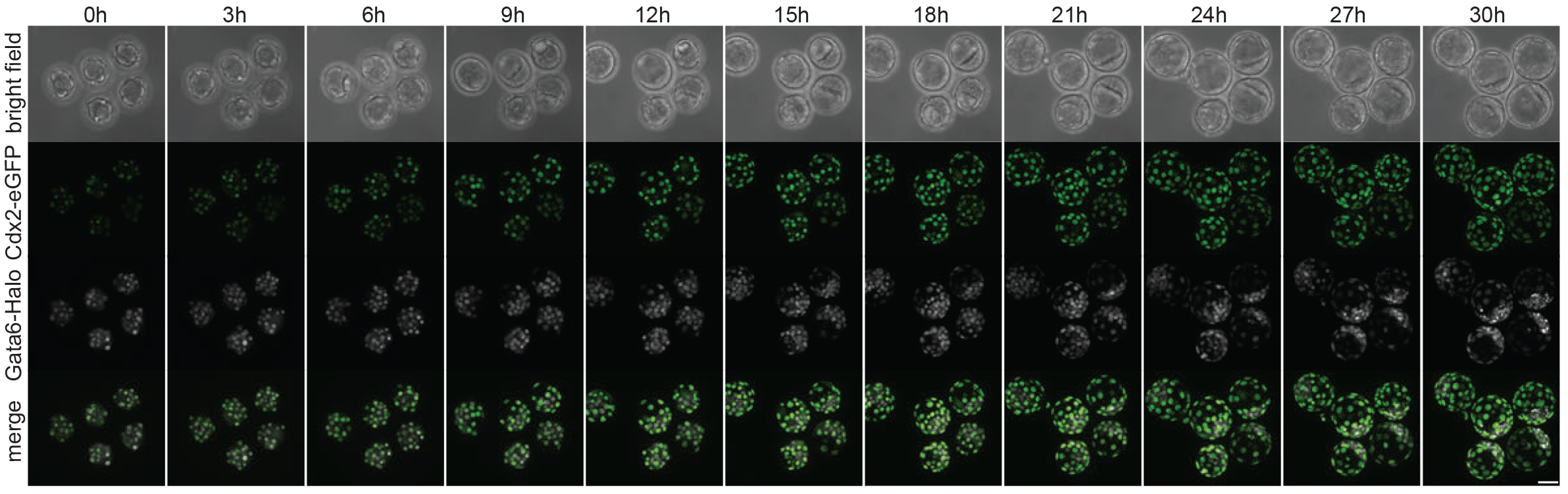
Application of multi-color reporter embryos in live imaging. Snapshots from time-lapse imaging of non-mosaic Cdx2-eGFP/Gata6-Halo embryos from the 16-cell stage to the blastocyst stage. The Halo-tag was visualized using the live JF549 dye. 1 Z-stack image was acquired every hour for 30 hours with a spinning disk confocal. All images show extended focus. Scale bars: 40 μm.

**Supplementary Figure 6.**
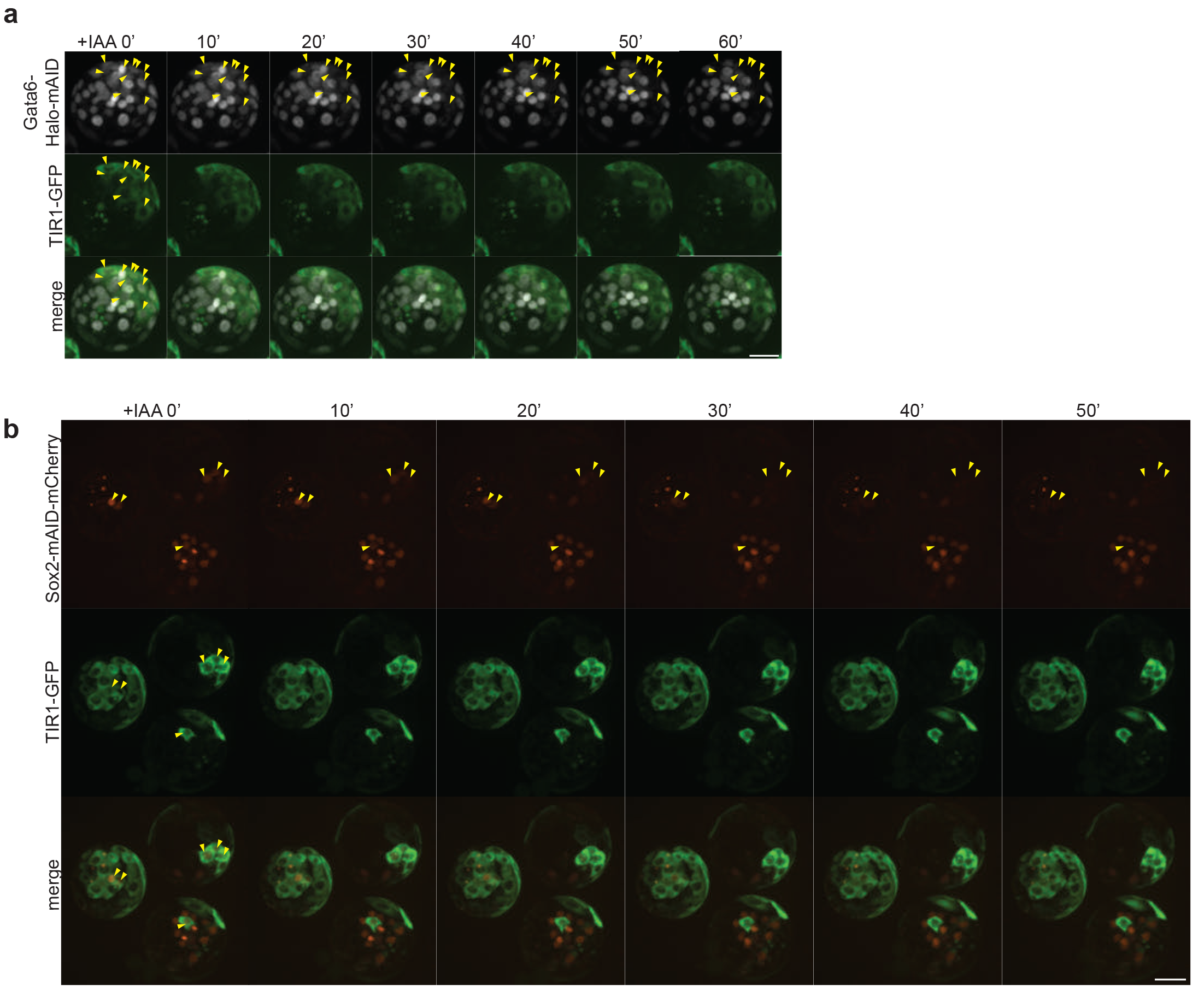
Examples of auxin-inducible degradation of Gata6-Halo-mAID and Sox2-mAID-mCherry in preimplantation embryos. Simultaneous targeting of Halo-mAID to the *Gata6* locus or mAID-mCherry to the *Sox2* locus and TIR1-GFP to the *Actb* locus was achieved with 2C-HR-CRISPR. Z-stack images were acquired every 10 minutes following addition of IAA. Arrows point to double positive cells with both Gata6-Halo-mAID or Sox2-mAID-mCherry, and TIR1-GFP expression. Extended focus images are shown. Scale bars: 40 μm.

## Supplementary Movie 1

Cytoplasmic microinjections of 2-cell stage mouse embryos.

## Supplementary Movie 2

Live imaging Cdx2-eGFP/Gata6-Halo embryos from the 16-cell to blastocyst stage.

